# Molecular dialogue between Orthonairovirus and tick: RNA-protein interactome of Hazara virus, a BSL2 model of Crimean-Congo Hemorrhagic Fever virus, in *Hyalomma* cells

**DOI:** 10.64898/2026.03.23.713610

**Authors:** Sarah Thibaudeau, Axel Grot, Alejandra Wu-Chuang, Yves Unterfinger, Véronique Legros, Margaux Ligner, Atéa Nermont, Lesley Bell-Sakyi, Houssam Attoui, John N. Barr, Roger Hewson, Guillaume Chevreux, Marion Sourisseau, Jennifer Richardson, Sandrine A. Lacour

## Abstract

Climate change and ecosystem collapse promote geographic expansion of vector-borne diseases, as witnessed by the recent incursions into Spain of the virus responsible for Crimean-Congo hemorrhagic fever (CCHFV). CCHFV is maintained in a tick-vertebrate cycle, principally involving ticks of the genus *Hyalomma*. Faced with the spread of *Hyalomma* ticks, and therefore the threat of a natural introduction of CCHFV into Western Europe, appropriate surveillance tools and control measures need to be implemented. It is both within and by the tick that CCHFV is maintained and spread in the environment. Despite prolonged portage of the virus, the tick is not overtly affected by CHFV infection. One of the prerequisites in conceiving control strategies is to understand the molecular mechanisms that intimately link the virus to its arthropod host. Despite the central role of the tick in the biology of CCHFV, these mechanisms are ill-defined, owing in part to the constraints associated with handling CCHFV-infected ticks in biosafety level 4 containment.

In this study, we established the network of interactions between the S segment of the RNA genome Hazara virus (HAZV), a BSL-2 model of CCHFV, and *Hyalomma* proteins using ChIRP-MS technique. We identified 166 tick proteins, 21 of which have been described as RNA-binding proteins. Gene ontology and pathway enrichment analyses revealed that the S segment RNA interacts predominantly with mitochondrial proteins that belong to various mitochondrial metabolic pathways.

## Introduction

Climate change and the collapse of ecosystems are leading to upheavals in the epidemiological patterns of infectious diseases, particularly vector-borne diseases. Vector-borne diseases, i.e. diseases whose causative agent is transmitted by a blood-feeding arthropod vector, have been a growing public health problem worldwide since the 21st century. Changes in the distribution of vector arthropod populations have led to the introduction of the pathogens they transmit into regions that were previously free of them (1). In this context, the World Health Organization established a global vector control strategy in 2017. (WHO, available on: [WHO EMRO - Global strategy for vector control](https://www.emro.who.int/fr/malaria-and-vector-borne-and-neglected-tropical-diseases/vector-control/global-strategy-for-vector-control.html)).

This phenomenon is clearly illustrated by the recent incursions into Western Europe of the virus responsible for Crimean-Congo hemorrhagic fever (CCHFV), principally transmitted by bites of ticks of the genus *Hyalomma.* In humans, CCHFV causes fever and hemorrhage, accompanied by a generally high mortality rate (10 to 40%)(2). Of note, CCHF has appeared since 2015 on the WHO’s list of emerging infectious diseases most likely to cause major epidemics.

The CCHFV genome was first evidenced in 2010 in *H. lusitanicum* ticks collected in south-western Spain (Estrada-Peña et al., 2012a) (3). In 2016, in the same region, a man died following multiple organ failure due to CCHFV and a nurse exposed to the patient’s blood became infected. The index patient had no history of travel and had probably been bitten by a tick (Negredo et al., 2017) (4). Since then, one indigenous case from 2013 has been retrospectively identified, and 17 other cases occurred between 2018 and 2025, including four additional deaths (5, Seasonal surveillance of Crimean-Congo haemorrhagic fever (CCHF) in the EU/EEA for 2025).

In France, although *H. marginatum* has been present in Corsica for several decades, it was first described in the south of France only in 2015 (6). This tick has very quickly spread throughout the Mediterranean region (7) and in October 2023, CCHFV was detected in *H. marginatum* ticks collected from cattle in the French Pyrénées-Orientales region (8).

CCHFV is maintained in a tick-vertebrate-tick cycle and is thought to spend 95% of its life cycle in ticks (9). After acquisition of CCHFV, the tick maintains the virus for its entire life span (several months to several years) and assures its transmission by not only horizontal but also vertical routes. Thus, in addition to its role as a vector, the tick serves as a reservoir for CCHFV (10). Despite the central role of the tick in the biology of CCHFV, very few studies have addressed the molecular mechanisms that confer the capacity of ticks to sustain prolonged CCHFV infection and ensure viral transmission (11, 12).

The persistence of vector-borne viruses is intimately linked to the life cycle of their arthropod vector: for ticks belonging to the *Ixodidae* family, the development cycle comprises 3 stages (larva, nymph et adult), the passage from one stage to the next requiring a moult that obligatorily follows a single bloodmeal. As the interval between two bloodmeals can be up to several months, the virus must be capable of not only surviving without substantial impairment of the life history traits of the tick, but also resisting successive moults. The organs in which the virus persists in the course of the different moults have not as yet been identified.

CCHFV belongs to the genus *Orthonairovirus*, family *Nairoviridae*, order *Hareavirales* and class *Bunyaviricetes* (13). Its viral genome consists of three single-stranded negative-polarity RNA segments. The S (small) segment encodes the NP nucleoprotein and the NSs protein, the M (medium) segment encodes the precursor of the GPC glycoproteins, which is then cleaved by furin and SKI-1 cellular proteases into the MLD, GP38, Gn, NSm and Gc proteins and lastly the L (large) segment encodes the RNA-dependent RNA polymerase RdRp (14).

Viruses are obligate intracellular life forms, whose composition is reduced to the bare essentials. Their survival therefore depends on the functions of the host cell, which they hijack for their own benefit, but also on the strategies they employ to evade the host cell’s immune system. For competent arthropod vectors, their life history traits seem to be little affected by viral infection, suggesting regulation—albeit without elimination—of viral replication. The cohabitation between the virus and the vector must therefore involve extensive molecular dialogue, and more particularly between viral (RNA and proteins) and cellular components, and notably cellular proteins. Understanding the molecular dialogue between ticks and viruses is thus an interesting avenue of investigation for elucidating the molecular mechanisms that underlie the vector competence of a given tick for a virus of interest.

Viral RNA, as both the genetic template for viral protein synthesis and genome replication and the principal target for sensors of innate antiviral pathways interacts with numerous RNA binding proteins (RBP) of host cell origin (15). In this study, we have identified *Hyalomma* proteins that interact with the S segment of HAZV, a BSL2 model for CCHFV. Unexpectedly, the viral RNA has been found to interact predominantly with mitochondrial proteins that belong to several mitochondrial metabolic pathways, and more particularly those related to energy production.

## Materials and methods

### Materials

#### Cell line and virus

The HAE/CTVM9 cell line came from the Tick Cell Biobank at the University of Liverpool. This cell line is derived from embryos of the *Hyalomma anatolicum* tick (16). The HAE/CTVM9 cell line was cultured at 30°C in L15/MEM medium (equivalent volume of L-15 (Leibovitz) (Gibco, Waltham, USA) and Minimal Essential Medium supplemented with Hank’s salts (Gibco, Waltham, USA) and 10% tryptose phosphate broth (Gibco, Waltham, USA), 20% fetal calf serum (Eurobio, Les Ulis, France), 2 mM L-glutamine (Gibco, Waltham, USA), 100 units/ml penicillin and 100 μg/ml streptomycin (Gibco, Waltham, USA).

The SW13 cell line (CCL-105, ATCC, Manassas, USA) is an epithelial cell line isolated in 1971 from the adrenal gland and cortex of a 55-year-old female patient with adrenal adenocarcinoma. SW13 cells were cultured in DMEM Glutamax medium (Gibco, Waltham, USA) supplemented with 10% FBS and 1% P/S.

Hazara virus (HAZV) strain JC280 (NCPV 0408084v, National Collection of Pathogenic Viruses, UK Health Security Agency, Salisbury, United Kingdom) was produced in SW13 cells.

#### Materials used for Mass-spectrometry analysis

MS grade Acetonitrile (ACN), H_2_O and formic acid (FA) were from ThermoFisher Scientific (Waltham, MA, USA). Sequencing-grade trypsin/Lys C mix was from Promega (Madison, WI, USA). Ammonium bicarbonate (NH_4_HCO_3_) was from Sigma-Aldrich (Saint-Louis, MO, USA).

### Methods

#### Infection kinetics

A total of 2.5 x 10^5^ HAE/CTVM9 cells were seeded in 24 wells plate and hermetically sealed 24h before infection with HAZV at an MOI of 0.1 or 0.01. At each collection time point, the cell pellets were collected for RNA extraction and quantification of the HAZV genome.

#### HAZV RT-qPCR

RNA extractions were performed using the NucleoSpin RNA Plus kit (Macherey-Nagel, Düren, Germany) for cell extractions, according to the manufacturer’s instructions. HAZV viral replication was then assessed in the HAE/CTVM9 cell line by RT-qPCR using TaqMan™ probe technology. Briefly, the extracted RNA was added to a mix consisting of the qScript™ XLT One-Step RT-qPCR ToughMix® (Quantabio, Beverly, MA, USA), specific HAZV primers (0.9µM, Appendix 1), and a HAZV probe (1.25µM, Appendix 1). The reaction volume was adjusted with water to a final volume of 10 µL. The following amplification program was used: 50°C for 10 min, 95°C for 1 min, then 45 cycles at 95°C for 10 s and 60°C for 45 s. The reaction was performed in a 96-well plate using the LightCycler 96 system (Roche, Basel, Switzerland). The copy number of transcripts was calculated using the standard curve method with a range of reference samples (HAZV segment S) and the results expressed as the number of RNA copies/ng of total RNA extracted (intracellular extraction). The standard was prepared by reverse transcription of a plasmid containing a DNA copy of the HAZV-S segment (17) by using the T7 Megascript RNAi kit according to the manufacturer’s instructions.

#### Cell infections

For HAZV infection, 30 x 10^6^ HAE/CTVM9 cells were cultured for 16h, then infected with HAZV at an MOI of 0.1. After 7 days, cells were collected and washed twice with PBS, followed by chemical crosslinking in a solution of PBS containing 4% formaldehyde at room temperature for 30min. Crosslinking was stopped by adding 2M glycine to a final concentration of 125mM and rotation for 5min at room temperature, followed by centrifugation at 330× *g* for 7 min. All supernatants were eliminated, and the final cell pellets were weighed. Cells were then frozen at −80°C until used. Non-crosslinked cells were processed the same way without the step of crosslinking, and mock-infected cells were cultured and handled in the same way as the infected and crosslinked cells, save that they were inoculated with a non-infectious inoculum (DMEM).

#### Comprehensive identification of RNA-binding proteins by mass spectrometry (ChIRP-MS)

HAZV targeting oligonucleotide probes were designed online using the Stellaris Probe designer tool (https://www.biosearchtech.com/stellaris), with repeat masking setting of 2 and even coverage of the whole S segment of HAZV. Full probe sequences are available in supplementary file 1 (probe sequences covering the whole segment S of HAZV used for ChIRP-MS.doc). Oligonucleotides were synthesized with a 3′ biotin-TEG modification by Eurofins Genomics (Luxembourg, Luxembourg).

ChIRP-MS was performed as previously described (18). Lysates were generated by resuspending cell pellets in 1 mL lysis buffer (50 mM Tris–HCl, pH 7.0, 10 mM EDTA, 1% SDS, supplemented with RNase and protease inhibitors) per 100 mg of cell pellet. Lysates were sonicated using a Hielscher UP200St sonicator (Teltow, Germany) with cycles of 30 seconds ON and 45 seconds OFF at 20W for 35 minutes. Then, 2mL of ChIRP hybridization buffer (750 mM NaCl, 1% SDS, 50 mM Tris-HCl pH 7.0, 1 mM EDTA, 15% formamide, supplemented with RNases and proteases inhibitors) was added per mL of sonicated lysate. Preclearing was achieved by adding 50µL of washed (following the manufacturer’s procedure) Dynabeads MyOne C1 (Invitrogen, Carlsbad, USA) per mL of lysate and rotation at 37°C for 30min. Preclearing beads were removed twice from the lysates using a magnetic stand; for this and all subsequent magnetic stand steps, 2min of separation were respected before removing any supernatant. To capture the vRNA-protein complexes, 1.5µL of 100µM ChIRP probe pool were added per mL of lysate. The ChIRP probe pool was composed of an equimolar mix of 10 antisense oligonucleotide probes. Hybridization was then performed by rotation at 37°C overnight. Afterwards, 150 µL of washed Dynabeads MyOne C1 per mL of lysate were added to the samples and incubated with rotation for 45 min at 37°C. Enriched material was collected on the beads using a magnetic stand, and beads were washed three times in 500µL ChIRP wash buffer (2x NaCl-sodium citrate (SSC buffer, Invitrogen), 0.5% SDS), each wash involving rotation for 10 min at 37°C and subsequent removal of supernatant using the magnetic stand. A second round of washing was performed 2 times with 500µL of a separate wash buffer (50 mM Tris-HCl, pH 7.0, 150 mM NaCl), following the same procedure. Finally, the beads were washed once with RNase-free water and resuspended in RNase-free water for MS processing. ChIRP-MS control experiments with mock-infected cells were performed identically by using the same set of probes.

#### Sample preparation prior to LC-MS/MS analysis

Beads from pulldown experiments were incubated overnight at 37°C with 20 μL of 50 mM NH_4_HCO_3_ buffer containing 1 µg of sequencing-grade trypsin/Lys C mix. Before LC-MS/MS analysis, the digested peptides were loaded and desalted on evotips purchased from Evosep (Odense, Denmark) according to the manufacturer’s procedure.

#### LC-MS/MS acquisition

Analyses were performed on a timsTOF Pro HT mass spectrometer (Bruker Daltonics, Bremen, Germany) coupled to an Evosep One system (Evosep, Odense, Denmark), operated with the manufacturer’s 40SPD Whisper Zoom method. This method relies on a 32.5-min gradient and a total cycle time of 36 min. Peptide separation was carried out on a C18 analytical column (0.075 × 150 mm, 1.7 µm beads, Aurora Elite CSI, IonOpticks), heated at 50 °C and operated at 200 nL/min. Mobile phases were (A) H₂O/0.1% FA and (B) ACN/0.1% FA. The timsTOF Pro HT was operated in DIA-PASEF mode, based on 12 pyDiAID frames, each with three mass windows, resulting in a 0.975 s cycle time as described in Bruker Application Note LCMS 218. Collision energy was applied stepwise as a function of ion mobility (19).

#### Data analysis

The raw MS files were processed using Spectronaut (version 20.3.251215.92449). A database search was performed against the *Hyalomma asiaticum* database from VectorBase (Hyas-2018, downloaded 01-2026, containing 29,644 entries), to which three additional sequences from the HAZV were added: NP, GPC, RdRp.

The search parameters included a dynamic calibration search and specific tryptic cleavage with up to two missed cleavages and three variable modifications. The considered modifications were acetylation (protein N-term), oxidation (M) and deamidation (N) as variable and Cys-Cys (C) as fixed.

Identifications were filtered using a 1% Q-value threshold at both the precursor and protein levels. Quantification was carried out using the Spectronaut Quantification Module at the MS2 level using areas. Protein inference was performed using the IDPicker algorithm.

#### Statistical Analysis

Multivariate statistics on protein measurements were performed using Qlucore Omics Explorer 3.9 (Qlucore AB, Lund, *SWEDEN*). A positive threshold value of 1 was specified to enable a log2 transformation of abundance data for normalization *i.e.* all abundance data values below the threshold will be replaced by 1 before transformation. The transformed data were finally used for statistical analysis *i.e.* evaluation of differentially present proteins between two groups using a Student’s bilateral t-test. A q-value better than 0.05 was used to filter differential candidates.

#### Identification of *H. asiaticum* orthologous proteins

Orthologous proteins were identified using a custom pipeline (Supplementary File 2: R_script_orthologs_identification.R) based on local BLAST searches implemented in R (4.4.3) within RStudio (2025.9.1.401), using local BLAST+ (version 2.17.0+) searches against publicly available proteome datasets for *Drosophila melanogaster* (UP000000803), *Homo sapiens* (UP000005640), *Ixodes scapularis* (UP000001555), and *H. asiaticum*.

Forward BLASTP searches were performed with an E-value cutoff of 1 × 10^-3^, retrieving up to 15 hits per query. Hits were post-processed to calculate adjusted sequence identity and coverage by merging overlapping alignments to better reflect true homology.

To identify inparalogs within the *H. asiaticum* proteome, an all-against-all BLASTP was run with stricter thresholds (E-value ≤ 1 × 10^-3^, identity ≥ 90%, coverage ≥ 80%). Inparalogous pairs were grouped into clusters using graph-based connectivity, accounting for recent duplications that could affect orthologue inference. This auto-BLAST script was implemented in R and is available in Supplementary File 3 (R_script_inparalogs_identification.R).

Reciprocal best hits (RBH) were determined by performing reverse BLASTP of hit sequences against the *Hyalomma* proteome. Orthology was assigned when query and subject proteins were mutual best hits or belonged to the same inparalog cluster, accommodating gene duplications. Final orthologous relationships were retained based on adjusted percentage identity ≥ 30%, adjusted query coverage ≥ 60%, and confirmation by RBH analysis with inparalog cluster adjustment.

#### Gene Ontology (GO) analyses

Gene Ontology (GO) enrichment analyses were performed to identify over-represented biological processes (BP), molecular functions (MF), and cellular components (CC) associated with orthologues from *I. scapularis*, *H. sapiens* and *D. melanogaster* (Supplementary File 4: goterm_supplementary).

GO term over-representation analysis was performed separately for each species using g:Profiler (g:GOSt) (20). Statistical significance of enrichment was evaluated using the cumulative hypergeometric test (one-sided Fisher’s exact test), comparing the query gene list against the default organism-specific background defined by g:Profiler (all annotated genes for the corresponding species). Multiple testing correction was applied using the Bonferroni method, and GO terms with an adjusted p-value < 0.05 were considered significant.

To reduce redundancy among enriched GO terms, semantic similarity-based clustering was performed on the combined set of enriched GO terms from the three species using REVIGO, with a similarity cutoff of 0.5 for BP and 0.7 for MF and CC. The resulting non-redundant GO terms were retained for downstream analyses and visualization.

Enrichment results were visualized using heatmaps based on −log10(Bonferroni-adjusted p-values) using GraphPad Prism version 10.

#### Pathway enrichment analysis

The pathway enrichment analysis was performed using the Reactome Pathway Analysis tool (Reactome version 95) from the list of human orthologuess (21). Over-representation analysis was conducted using a hypergeometric test and p-values were adjusted for multiple testing using the Benjamini–Hochberg false discovery rate (FDR) correction. Pathways with an FDR-adjusted p-value ≤ 0.05 were considered significantly enriched

## Results

### Intracellular HAZV RNA peaked at 7 days after infection

The prerequisite for establishing the interactome of the S segment of HAZV was to define the conditions of infection for which the intracellular amount of viral RNA is maximal. We therefore monitored intracellular viral replication over time (fig 1). The amount of intracellular viral RNA increased by 4 logs between the day of infection and day 7, remained stable over the following days, and then began to decrease slightly between days 13 and 15 post-infection.

**Figure 1.**
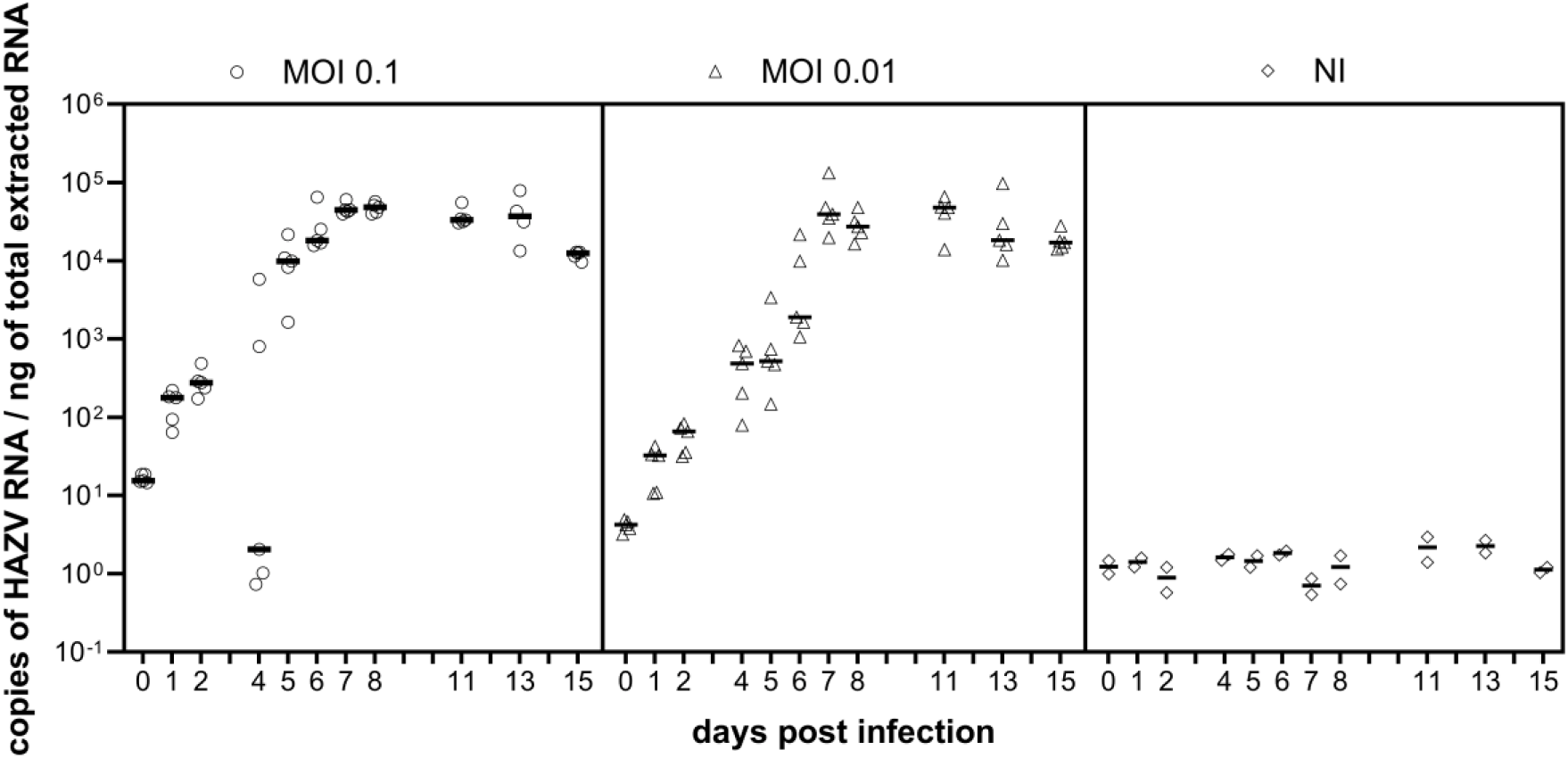
Replication HAZV in HAE/CTVM9 tick cells. HAE/CTVM9 tick cells were infected with HAZV at an MOI 0.1 (○) or 0.01 (Δ) or were not infected (◊). Viral RNA from cells of five replicates per day was quantified by RTqPCR and expressed as copies of HAZV RNA/ng of total extracted RNA.

Infection was subsequently carried out on a total of 30.10^6^ cells at an MOI of 0.1 for 7 days. At the end of these 7 days and prior to processing for ChIRP analysis, successful infection of cells was verified by quantification of HAZV RNA (data not shown).

It should be noted that the values of the five replicates per day are relatively consistent, except for day 4 of infection at MOI 0.1, for which three replicates appear not to have been infected.

### The tick host protein interactome of the S segment of HAZV was defined using ChIRP-MS

Comprehensive Identification of RNA binding Protein by Mass Spectrometry (ChIRP-MS) is an RNA-centric, unbiased approach that can specifically identify all proteins interacting with a given RNA sequence, here the S segment of HAZV, therefore referred to protein complexes, some which are RBPs (RNA-Binding Proteins). This technique has the advantage of being highly sensitive due to the use of mass spectrometry.

Technically, ChIRP-MS can be broken down into five main steps: (i) crosslinking of RNA-protein complexes, (ii) hybridization between probes and viral RNA, (iii) oligoprecipitation, of RNA-protein complexes (iv) elution of protein complexes, and (v) protein identification by mass spectrometry. The methodology of this approach is summarized in figure 2.

**Figure 2.**
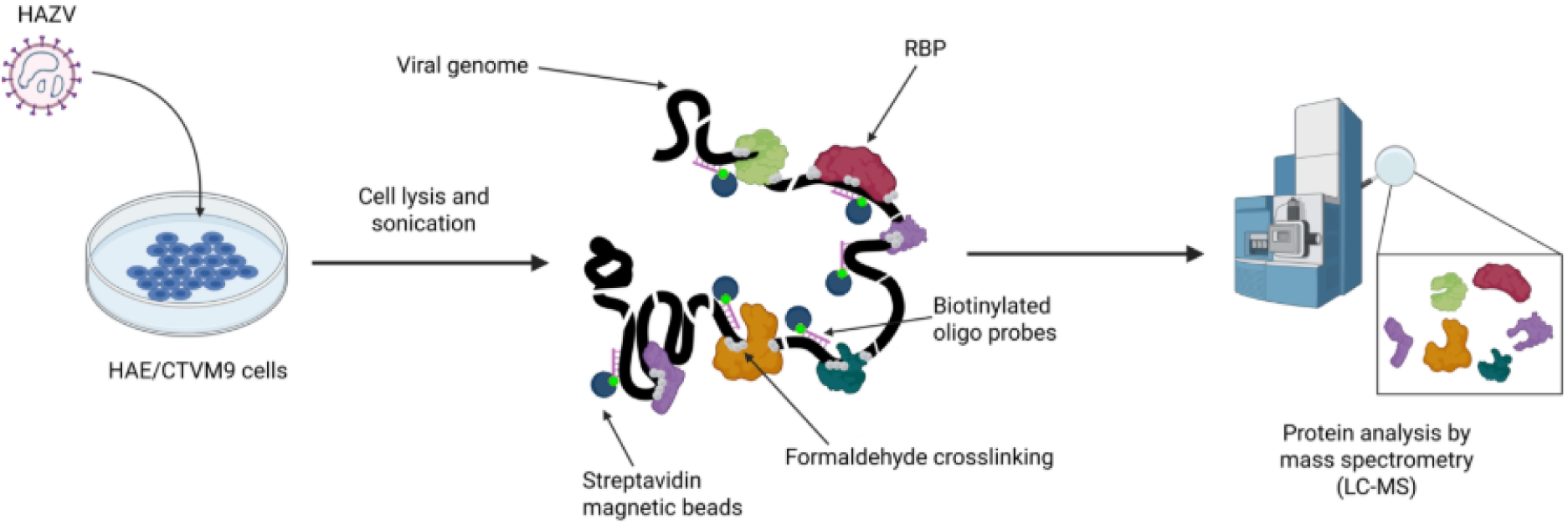
Strategy for identifying tick proteins interacting with HAZV RNA by ChIRP-MS. Briefly, after infection of cells, RNA-protein molecular complexes are hybridized with biotinylated antisense oligonucleotides. Viral RNA and associated protein complexes are next captured and purified using streptavidin-coupled beads. The co-precipitated proteins are then identified by LC-MS/MS.

ChIRP-MS was performed on four infected replicates and four uninfected replicates. One replicate from each condition was excluded due to disparity compared with the other three (data not shown).

### ChIRP-MS and Blast searches in the *H. asiaticum* proteome identified 166 tick proteins that interact with the S segment of the HAZV

We identified 169 proteins (including the three viral proteins that were sought) with the parameters of fold change ≥ 2 and adjusted q-value ≤ 0.05, that we define as the enlarged interactome (figure 3). When we adjust the parameters of the q-value to ≤ 0.01, this list is shortened to 32 proteins, including three viral proteins, and is defined as the high-confidence, restricted interactome. These viral proteins were the most enriched, in keeping with the known interactions between the viral genome and the viral proteins NP and RdRp, which all together constitute the viral ribonucleoprotein complex.

**Figure 3.**
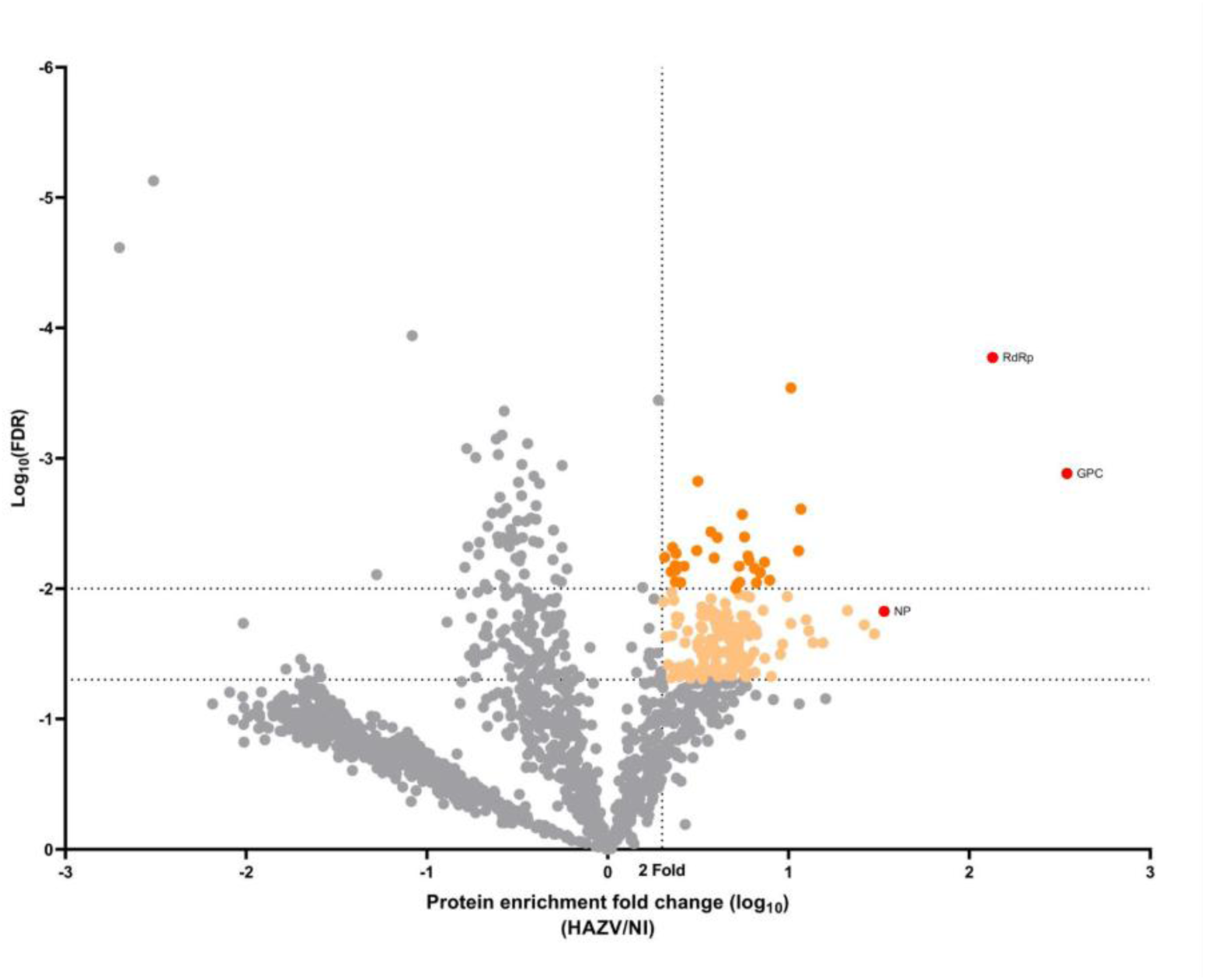
Representation of proteins enriched in the S segment interactome of HAZV RNA. The volcano plot displays the log_10_ fold change (x-axis) versus the −log₁₀ adjusted p-value (y-axis) for all genes detected in the dataset. Each point represents a gene. Significantly enriched genes in the infected condition compared with the mock condition (|log₂FC| ≥ 1; q-value ≤ 0.05) are highlighted in salmon or (|log₂FC| ≥ 1; q-value ≤ 0.01) in orange. The proteins of HAZV are represented in red. Vertical and horizontal dashed lines indicate the significance thresholds.

The 166 proteins of *H. anatolicum* identified with reference to the proteome of *H. asiaticum* were arbitrarily named Hyan1 to Hyan166.

Due to the limited information available for ticks of the genus *Hyalomma*, which prevented us from conducting more in-depth analyses, we searched for orthologous proteins in three species: *I. scapularis*, *H. sapiens*, and *D. melanogaster*. For the 166 proteins of *H. anatolicum*, we identified 113 orthologues in *I. scapularis*, 125 in humans, and 118 in fruit flies (ChIRP_orthologues_hyalomma_filtered_adjusted_RBH_inparalogues_qvalue.xlsx). Of note, for 34 *Hyalomma* proteins orthologues were not found in any of the three species.

ChIRP-MS is a technique designed to capture RNA-binding proteins, but in reality, this technique can identify all proteins complexed with RBPs. Using the list of human orthologues, we were able to identify 21 proteins already described as RBPs, and 6 for which the RBP function has not been demonstrated but is suspected (table 1).

**Table 1.** Identification of RNA-binding proteins among human orthologs.

| Hyan_ID | human_ID | protein name | RBP status | presumed RBP function |
| --- | --- | --- | --- | --- |
| Hyan57 | Q9UI43 | rRNA methyltransferase 2, mitochondrial (MRM2) | RBP confirmed | Mitochondrial rRNA modifying enzyme; direct binding to rRNA |
| HyAn23 | Q7L0Y3 | tRNA methyltransferase 10 homolog C (TRMT10C / MRPP1) | RBP confirmed | Mitochondrial tRNA modifying enzyme |
| HyAn132 | Q9BYD1 | Large ribosomal subunit protein uL13m (MRPL13) | RBP confirmed | Mitochondrial ribosomal protein; direct interaction with rRNA |
| HyAn61 | Q9BSH4 | Translational activator of cytochrome c oxidase 1 (TACO1) | putative RBP | Mitochondrial translocation factor; functional interaction with mt-COX1 mRNA |
| HyAn18 | A4D1E9 | GTP-binding protein 10 (GTPBP10) | RBP confirmed | Mitochondrial ribosome assembly factor; direct interaction with rRNA |
| HyAn66 | E7ETU7 | Large ribosomal subunit protein uL3m | RBP confirmed | Mitochondrial ribosomal protein; intrinsic rRNA binding |
| HyAn48 | Q07021 | Complement component 1 Q subcomponent-binding protein (C1QBP/p32) | putative RBP | Multifunctional protein; reported RNA interactions (rRNA/viral RNA) but no evidence and no canonical RBD |
| HyAn73 | P62241 | Small ribosomal subunit protein eS8 (RPS8) | RBP confirmed | Cytosolic ribosomal protein; direct interaction with rRNA |
| HyAn151 | Q96A35 | Large ribosomal subunit protein uL24m | RBP confirmed | Mitochondrial ribosomal protein; rRNA interaction |
| HyAn25 | Q13825 | Methylglutaconyl-CoA hydratase (AUH) | putative RBP | AUH is an enol-hydratase also known as AUH RBP (binding to mRNA AREs). |
| HyAn137 | Q9Y6V7 | ATP-dependent RNA helicase DDX49 | RBP confirmed | DEAD-box helicase; direct RNA binding demonstrated |
| HyAn36 | Q8IYB8 | ATP-dependent RNA helicase SUPV3L1 | RBP confirmed | Mitochondrial RNA helicase involved in mtRNA surveillance/degradation |
| HyAn103 | A0A804HJZ1 | Leucine-rich PPR motif-containing protein, mitochondrial | RBP confirmed | PPR domain, direct binding to mitochondrial RNAs (stability/translation) |
| HyAn112 | Q7L3T8 | Probable proline--tRNA ligase, mitochondrial (PARS2) | RBP confirmed | Aminoacyl-tRNA synthetase; direct binding of tRNA |
| HyAn139 | Q9H9Y2 | Ribosome production factor 1 (RPF1) | putative RBP | Ribosomal biogenesis factor; functional interaction with rRNA/pre-ribosomal complexes without direct CLIP evidence |
| HyAn20 | Q9NP81 | Serine--tRNA ligase, mitochondrial (SARS2) | RBP confirmed | Aminoacyl-tRNA synthetase; direct binding to tRNA |
| HyAn80 | Q9H9J2 | Large ribosomal subunit protein mL44 (MRPL44) | RBP confirmed | Mitochondrial ribosomal protein; intrinsic rRNA binding |
| HyAn136 | Q02543 | Large ribosomal subunit protein eL20 (RPL20) | RBP confirmed | Cytosolic ribosomal protein; rRNA interaction |
| HyAn142 | B0QYK0 | RNA-binding protein EWS (EWSR1) | RBP confirmed | Alternative isoform/entry of EWSR1; same RBP function |
| HyAn124 | Q15269 | Periodic tryptophan protein 2 homolog (WDR12) | putative RBP | Ribosomal biogenesis factor (PeBoW); functional interaction with pre-rRNA without direct CLIP evidence |
| HyAn140 | A0A6Q8PGZ8 | Glycine--tRNA ligase (GARS) | RBP confirmed | Aminoacyl-tRNA synthetase; direct binding to essential tRNA |
| HyAn115 | Q9Y2Q9 | Small ribosomal subunit protein bS1m (MRPS1) | RBP confirmed | Mitochondrial ribosomal protein; intrinsic rRNA interaction |
| HyAn121 | P43897 | Elongation factor Ts, mitochondrial (TSFM) | putative RBP | Mitochondrial translation elongation factor; functional interaction with mRNA/ribosome |
| HyAn56 | G3V325 | Pentatricopeptide repeat-containing protein 1, mitochondrial | RBP confirmed | PPR domain: direct binding to mitochondrial RNAs |
| HyAn105 | K7ES61 | Large ribosomal subunit protein uL4m (fragment) | RBP confirmed | Mitochondrial ribosomal protein; rRNA binding |
| HyAn17 | O95363 | Phenylalanine--tRNA ligase, mitochondrial (FARS2) | RBP confirmed | Aminoacyl-tRNA synthetase; direct binding to tRNA |
| HyAn164 | Q9P015 | Large ribosomal subunit protein uL15m (MRPL15) | RBP confirmed | Mitochondrial ribosomal protein; intrinsic rRNA binding |

This table summarizes the identification of RNA-binding proteins (RBPs) within a dataset of human orthologous proteins. Protein annotations were retrieved from UniProt, and each protein was categorized as a confirmed or putative RBP according to UniProt annotations, based on the presence of known RNA-binding domains, experimentally validated RNA interactions, or functional descriptions implying RNA-binding capacity.

### The segment S of HAZV is predominantly associated with mitochondrial proteins

To gain insight into the biological functions of the proteins interacting with the S-HAZV RNA, we performed Gene Ontology (GO) enrichment analysis based on the list of orthologues from *I. scapularis*, *H. sapiens*, and *D. melanogaster* (Figure 4).

**Figure 4.**
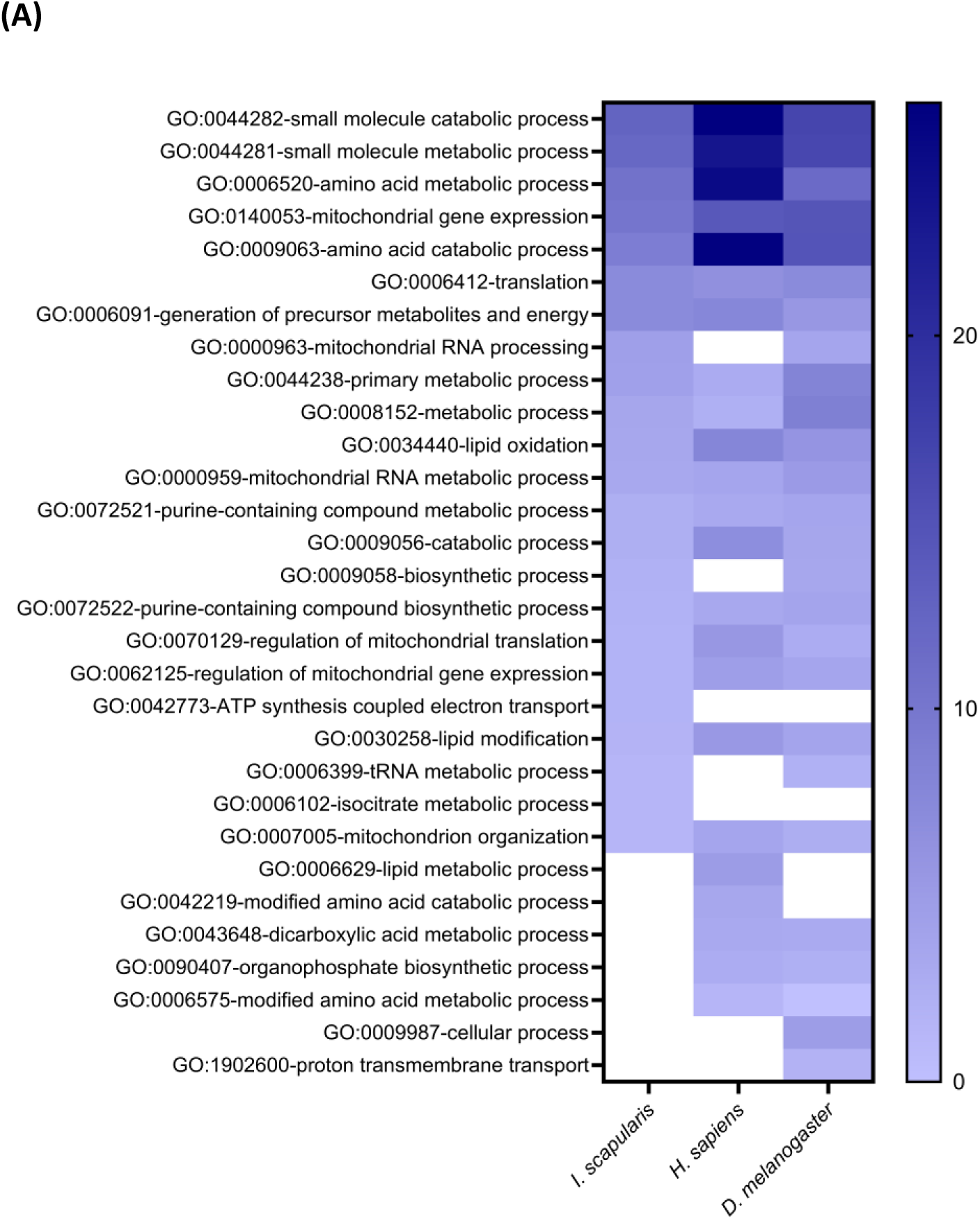
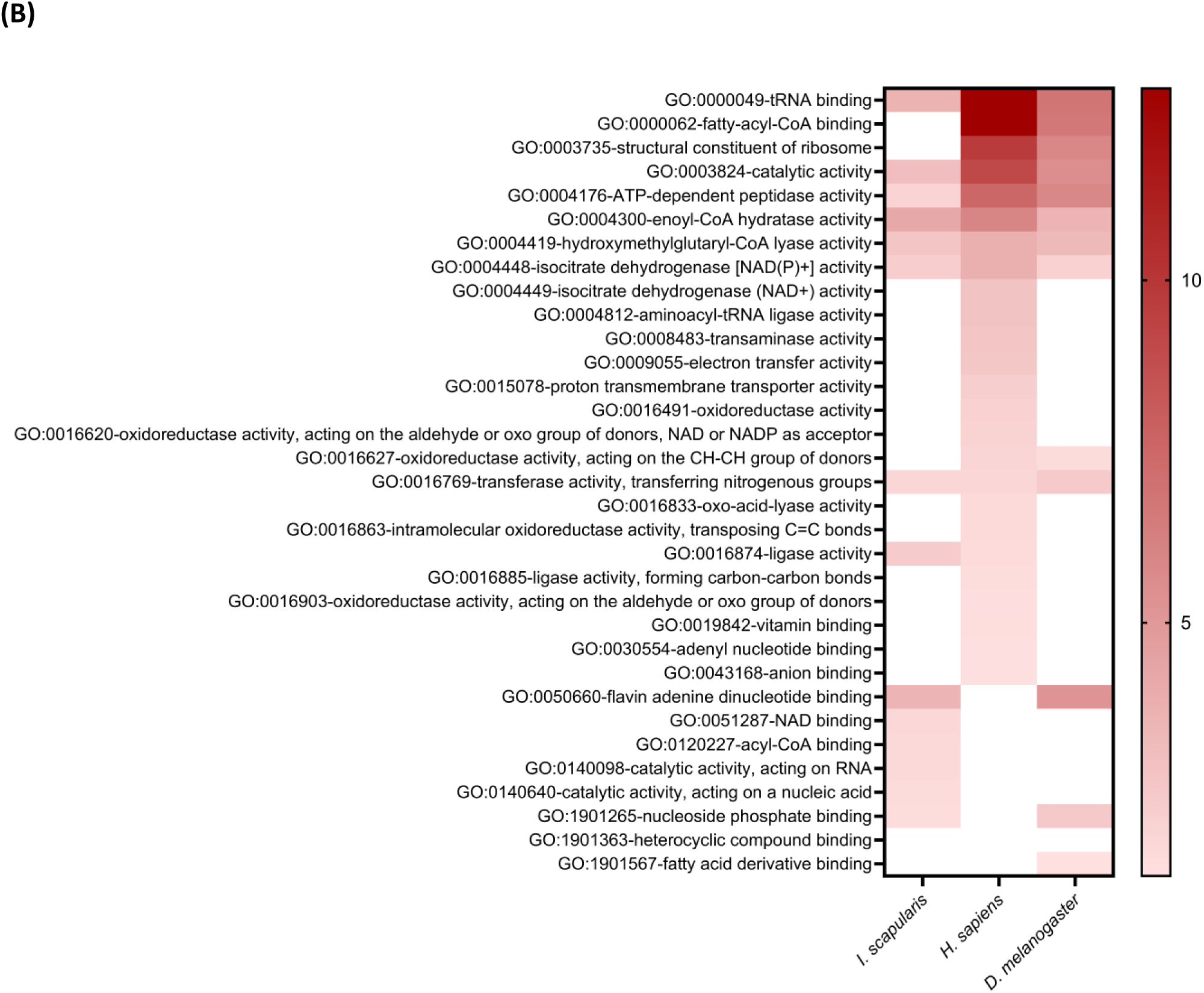
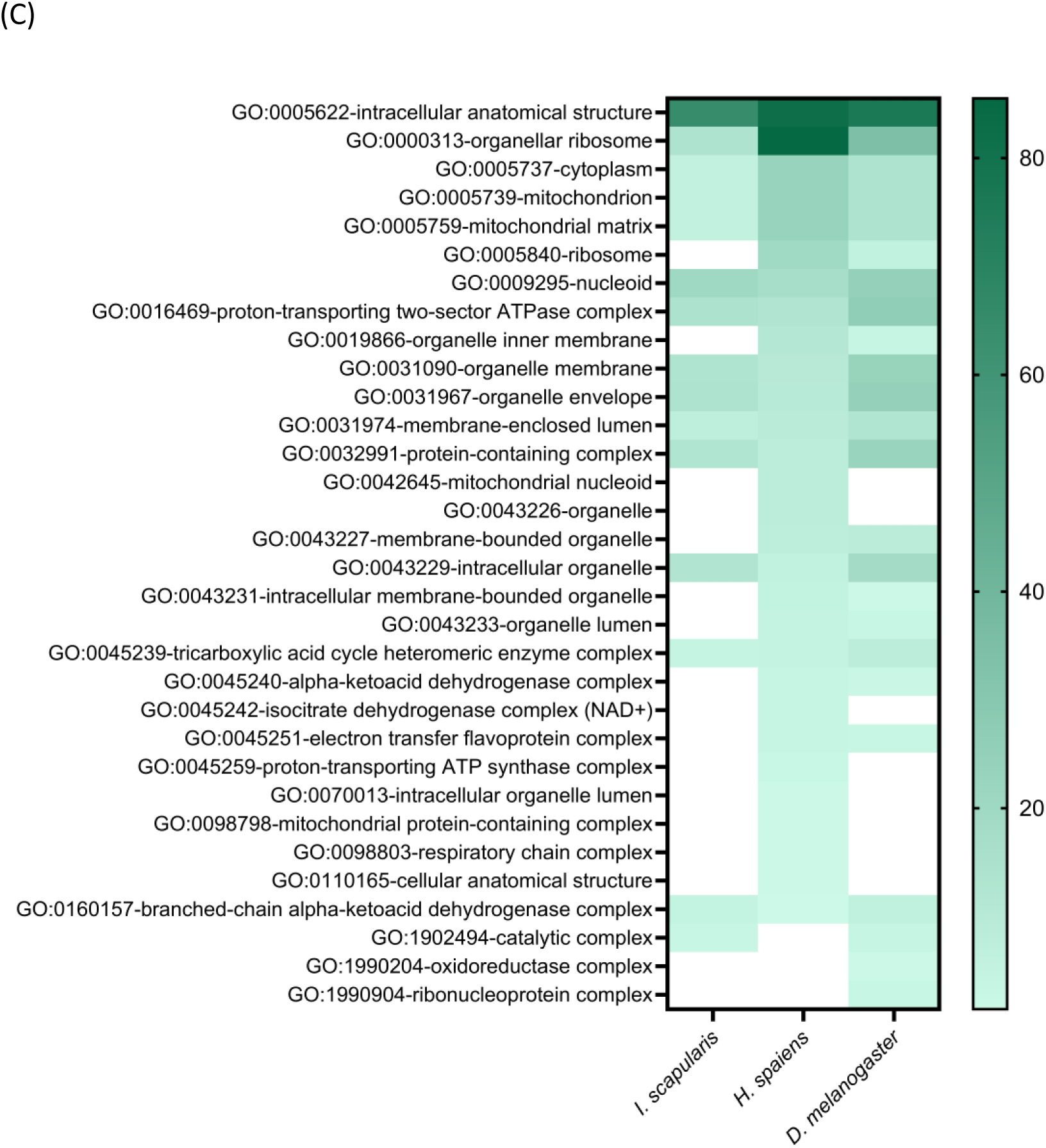
Comparative Gene Ontology (GO) enrichment analysis of orthologous proteins across three species. Heatmaps illustrate the enrichment of Gene Ontology (GO) terms associated with orthologous proteins from *Ixodes scapularis*, Homo sapiens, and *Drosophila melanogaster*. Panels represent the three GO categories: (A) Biological Process (BP), (B) Molecular Function (MF), and (C) Cellular Component (CC). GO enrichment analysis was performed using g:Profiler, applying a hypergeometric test (one-sided Fisher’s exact test). GO terms significantly enriched in at least one species were retained and subsequently reduced using REVIGO to minimize semantic redundancy (similarity threshold = 0.5 for BP; 0.7 for MF and CC), resulting in approximately 30 GO terms per heatmap. Heatmap values correspond to −log₁₀(Bonferroni-adjusted p-values), computed and visualized using GraphPad Prism version 10. Only GO terms meeting the significance threshold (adjusted p-value < 0.05) were considered. GO terms not significantly enriched for a given species are shown as white cells.

Across the three species, GO analyses of Biological Process, Molecular Function and Cellular Component revealed highly coherent enrichment patterns, with metabolic and biosynthetic processes supported by enzymatic molecular functions and localized predominantly to cytosolic and mitochondrial compartments, consistent with conserved cellular core functions. Several mitochondrial-related functions were significantly enriched, suggesting a possible involvement of mitochondrial pathways in the infection. Among other enriched biological processes, some were related to RNA metabolism, as expected. Overall, these results suggest that the S-HAZV RNA preferentially associates with proteins involved in multiple cellular pathways, those related to RNA, catabolic processes, or pertaining to mitochondrial functions being particularly prominent. The predominance of the association of *H. anatolicum* proteins with mitochondrial proteins can be appreciated in the representation of the interactome of the segment S of the HAZV, as drawn up with Cytoscape (22) (Figure 5).

**Figure 5.**
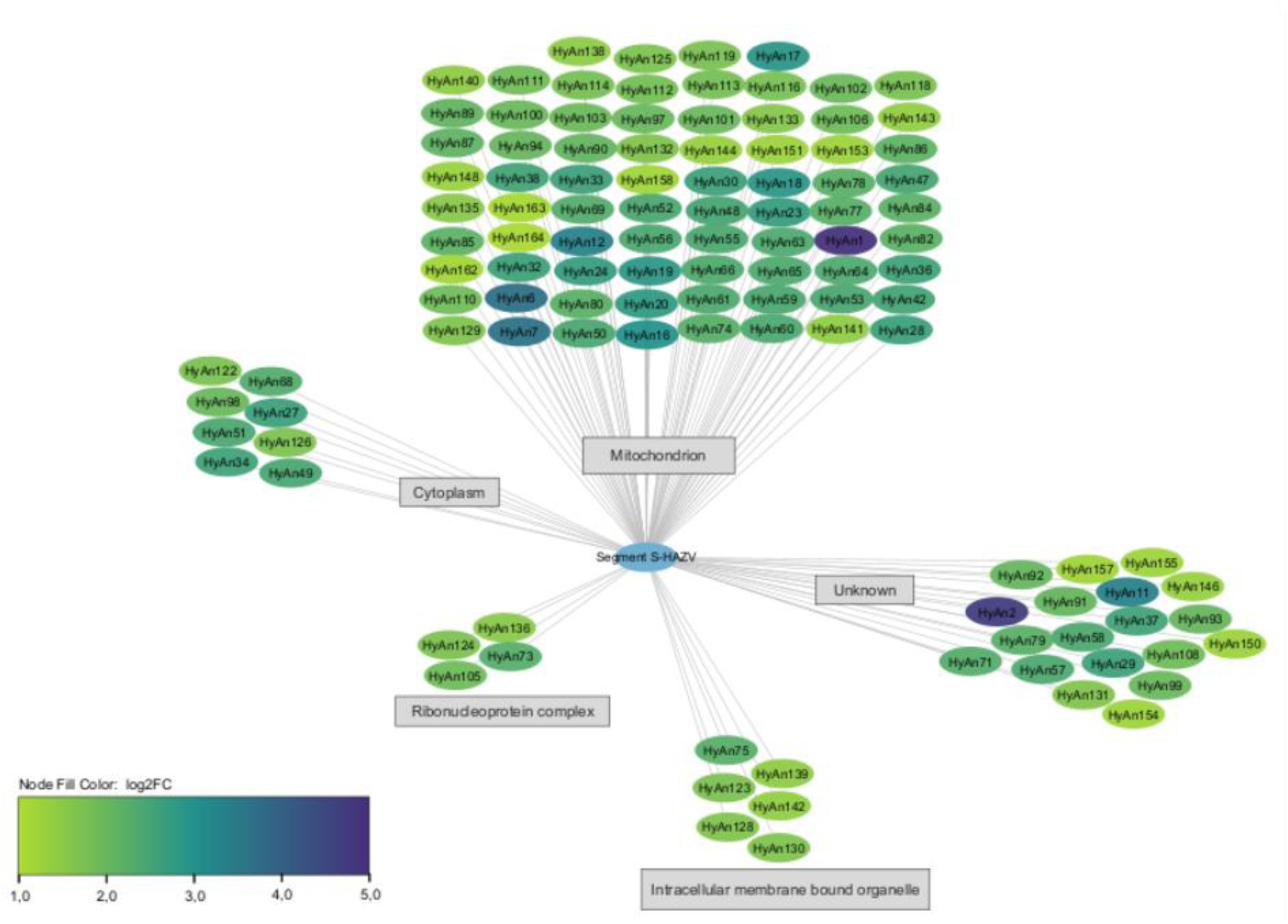
Cytoscape network of the S-HAZV–associated tick proteome in HAE/CTVM9 cells, inferred using the *Hyalomma asiaticum* proteome. Nodes represent proteins identified in the ChIRP experiment. Node color indicates ChIRP enrichment in HAE/CTVM9 cells at 7 days post-infection. Proteins were annotated and grouped according to their cellular compartment using g:Profiler by orthology with *Ixodes scapularis.* Proteins that do not have orthologs in *Ixodes scapularis* are not represented. The representation was made with Cytoscape 3.10.3.

### The segment S of the HAZV is mainly associated with proteins involved in mitochondrial pathways

Enrichment analysis of pathways via Reactome also showed that the majority of *Hyalomma* proteins that interact with the S segment of HAZV are involved in mitochondrial metabolic pathways, particularly in energy production pathways (respiratory chain, fatty acid beta-oxidation, ketone body metabolism (table 2).

**Table 2.**
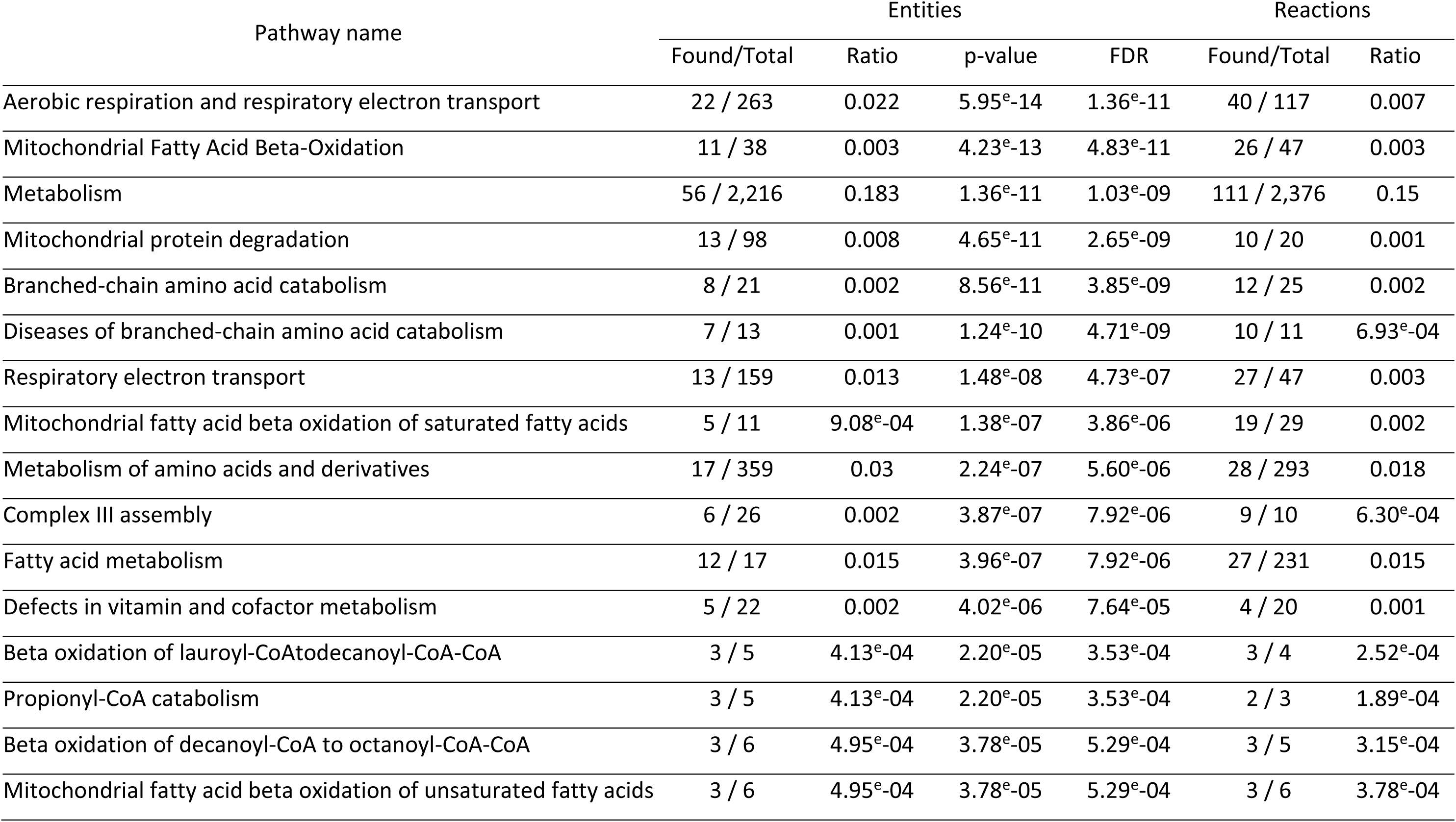
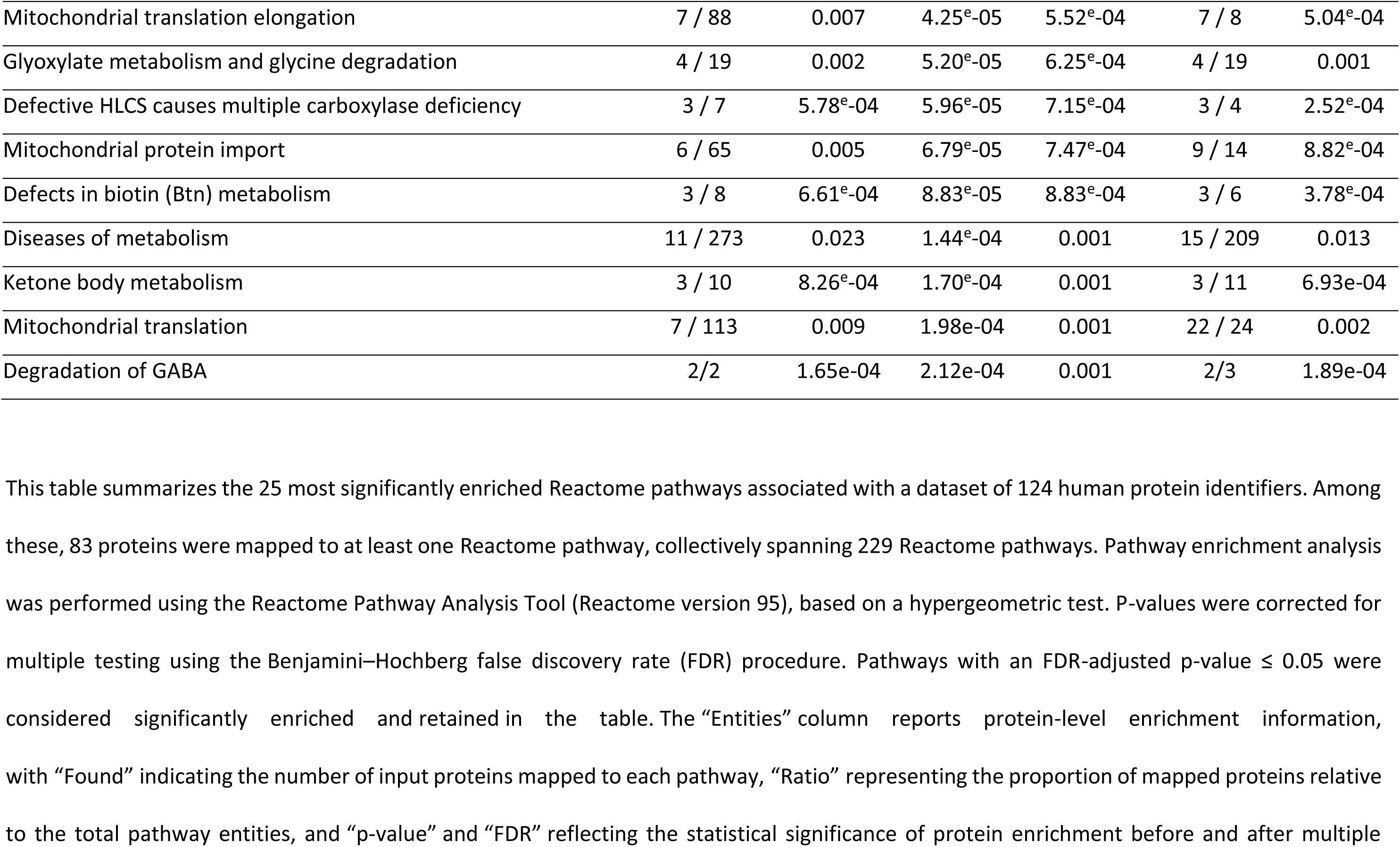
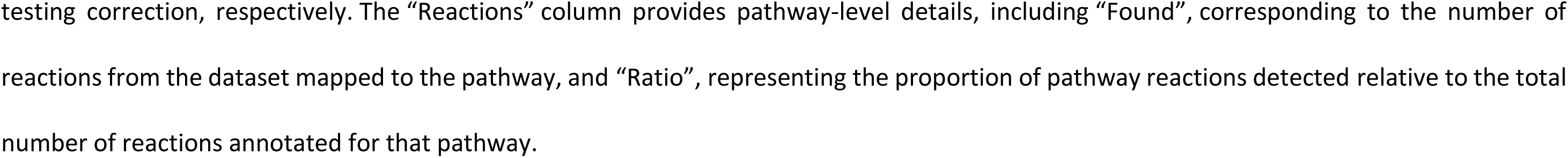
Reactome pathway enrichment analysis of human orthologous proteins.

## Discussion

Ticks represent not only vectors but also essential reservoirs for the orthonairovirus CCHFV. The molecular means by which the virus subverts biological processes and escapes elimination by tick immunity are largely unknown. Using a cell-based model of CCHFV infection of ticks — namely, infection of a *Hyalomma* tick cell line with the closely related HAZV — we have resolved the viral RNA interactome between the S segment of the HAZV genome and the proteome of a *H. anatolicum* cell line. Enrichment analysis of the set of RNA-associated tick proteins revealed an unexpected preponderance of proteins of mitochondrial origin. The significance of these findings can only be inferred with reference to current understanding of the molecular biology of genomic RNA of orthonairoviruses within infected cells.

The negative-stranded RNA genome of orthonairoviruses is a template for both viral transcription and genome replication (23). Viral transcription is primed by the capture of 5’ termini of capped cellular RNA, called “cap-snatching”, such that viral mRNA are capped and bear heterologous 5’ sequences of host cell origin (24). The genome is also copied into a positive-stranded antigenome, which then serves as the template for amplification of genomic RNA. Both transcription and genome replication are carried out within ribonucleoprotein (RNP) complexes, comprising both viral RNA and the viral structural proteins N and L, the latter of which both bind to viral RNA. The N protein envelops the viral RNA, while the L protein catalyzes cap-snatching, transcription and genome replication.

While for orthonairoviruses the processes of viral transcription and genome replication are fairly well understood at the molecular level, their relationship to cellular physiology is as yet ill-described. In particular, while viral transcription and genome replication are known to take place in the cytoplasm, their precise intracellular location is unknown. In a comparative study, Xu *et al.* (25) investigated the localization of ectopically expressed N protein – which has been considered a proxy for viral transcription and genome replication, due to its tight association with the viral genome – for several viruses of the class *Bunyaviricetes*, including CCHFV. For all examined viruses, the N protein was found to reside in major cytoplasmic RNA granules and, more particularly, in P bodies and stress granules (SG). As SG contain intact mRNAs that are simply stalled in translation, the authors noted that SG might providing an ideal location for cap-snatching. Furthermore, the N protein of a plant bunyavirus co-localized not only with the forementioned RNA granules, but also with a nucleocytoplasmic transport factor in the perinuclear region, raising the possibility that the viral RNP might capture the capped 5’ extremities of cellular mRNA as they enter the cytoplasm through nuclear pore complexes (25). This last finding is in accordance with the study of Andersson *et al.* (26), in which the N protein of CCHFV was localized to the perinuclear region of mammalian cells, whether in infected cells or upon ectopic expression of N protein, in relationship to the capacity of N to bind to actin filaments.

Our ChIRP-MS strategy was designed to retrieve protein complexes associated with the S segment of the HAZV genome. However, as genomic RNA is thought to be enclosed by the viral N protein, the accessibility of viral RNA to the complimentary oligonucleotide probes might be expected to be restricted. Nevertheless, genomic viral RNA may be more accessible than generally appreciated. In this regard, the N protein must be temporarily stripped from viral RNA for occupation of the active site of the L protein during transcription and genome amplification (23). Moreover, the ambisense S segment of the Rift Valley fever virus, a bunyavirus of the *Phenuiviridae* family, contains an exposed element that overlaps with the intergenic region between the N and NSs coding regions (27). The exposed region assumes a secondary structure that would appear to exclude N binding. Whether an analogous element exists in the S segment of orthonairoviruses is unknown. Finally, it has also been speculated that within viral factories, genomic RNA might not be enclosed by N protein to the extent transposed from *in vitro* studies (23). Binding of RBP to viral genomic RNA might be subject to similar limitations. However, since the ChIRP-MS strategy retrieves protein complexes associated with the targeted RNA and not solely RNA-binding proteins, only a fraction of the recovered proteins would be expected to interact directly to the S segment RNA. Since N and L proteins necessarily bind directly to genomic viral RNA, it is probable that at least some of the recovered host cell proteins interact with the N or L proteins, rather than with viral RNA.

The interactome of the S segment was expected to contain viral RNA-binding proteins N and L, as well as host cell proteins. The latter would include cellular proteins whose activity is subverted to support viral transcription and genome replication, and possibly host cell proteins that behave as sentinels or as effector proteins in antiviral defense pathways. As expected, the interactome of the S segment was indeed enriched for the expected viral proteins, N and L, as well as host cell proteins of considerable functional diversity.

Among the RBP found to be associated with the S segment of the HAZV genome, we expected that we might find sentinel or effector proteins belonging to arthropod innate antiviral pathways, and more particularly the RNAi pathway. The RNA genome of orthonairoviruses comprises double-stranded elements — such as the panhandles formed by complementary sequences at the extremities of the genomic segment, whose hybridization mediates circularization — that ought to be recognized by the Dicer protein. In this regard, ectopic expression of the S segment of HAZV in a tick cell line led to production of small interfering RNA and induction of HAZV S segment-specific RNA interference (12). Nonetheless, canonical components of the RNAi pathway, or of other antimicrobial pathways, were not identified within the S segment interactome. Several explanations can be advanced for the apparent absence of antiviral pathway proteins. First, constitutive expression of such proteins might be low, and if expression is transiently induced upon viral infection, the examined time-point (7 dpi) might have succeeded the rise in expression. Moreover, binding of sentinel proteins to viral RNA is expected to be impeded by the N protein: the quantity of bound Dicer might thus lie beneath our threshold of detection. Lastly, multiple RBP, some of which uncharacterized, were retrieved in association with S segment RNA: one or more of these might exert an unsuspected antiviral function.

Unexpectedly, a majority of the retrieved proteins were of mitochondrial origin. These proteins were derived from all mitochondrial compartments; that is, outer membrane, inner membrane and matrix and from multiple mitochondrial protein complexes. The connexion between mitochondrial proteins and HAZV RNA appears not to be specific to the tick cell context, as we made similar findings in a preliminary study performed in a human cell line. Of note, in our hands mitochondrial proteins did not predominate in the viral RNA interactome of a tick-borne orthoflavivirus, whether in a human or a tick cell line, as resolved by ChIRP-MS methodology (28) and manuscript in preparation). Our results thus suggest that the connexion between the viral genome — or at least its S segment — and mitochondrial proteins depends upon the virus rather than the host cell species. Whether the connexion might be shared by other orthonairoviruses, or even among *Bunyaviricetes*, is as yet unclear. To the best of our knowledge, no other description of the viral RNA interactome of *Bunyaviricetes* exists in the literature, although the impact of the Severe Fever with Thrombocytopenia Syndrome and Uukuniemi viruses (SFTSV and UUKV, respectively) on the global set of host cell RBP was resolved in tick cell lines (29, 30). Nevertheless, the interactome of the N protein with the tick cell proteomes was resolved for SFTSV and UUKV in the cited studies; while mitochondrial proteins were not preponderant in the N interactome, the extent of overlap between N and viral RNA interactomes is unknown; that is, complexes of mitochondrial proteins might be bound to viral RNA but not via the N protein.

Viral survival and mitochondrial function are intricately connected. Mitochondria house the molecular machinery of oxidative metabolism as well as biosynthetic and signalling pathways required by the host cell. Moreover, the intrinsic apoptotic pathway, which can be of benefit or detriment to the virus, depending notably upon the stage of the viral cycle (31), is triggered at the outer mitochondrial membrane. In mammalian cells, the outer membrane is also a platform for the induction of antiviral response, in that it houses the central adaptor protein mitochondrial antiviral signaling protein (MAVS), which lies downstream of viral RNA sensing by RIG-I-like receptors. Orthologues of RIG-I-like receptors and MAVS have not been identified in arthropods, and a role for mitochondria in the antiviral response has not been described. In mammalian cells, many viruses have been shown to impinge upon mitochondrial function, such as by inducing mitochondrial fission or fusion, by triggering mitophagy or by interfering with the function of MAVS. In many cases such interference has been interpreted as antagonizing the antiviral response (31). In the N protein of SFTSV interactome resolved by Petit et *al.* (29), mitochondrial stress factors and proteins associated with stress granules were found, and among these latter the Up-frameshift protein (UPF1), a component of the nonsense-mediated mRNA decay (NMD) pathway.

How and where genomic RNA of HAZV encounters mitochondrial proteins is currently unclear. To the best of our knowledge, RNA of orthonairoviruses — or at least its proxy, the N protein — has never been localized within or in proximity to mitochondria, suggesting that contact between viral RNA and mitochondrial proteins occurs outside mitochondria. The large majority of mitochondrial proteins are encoded by nuclear DNA and are imported into mitochondria subsequent to translation in the cytoplasm. Human mitochondrial DNA encodes thirteen mitochondrial proteins, which are translated within mitochondria. The mitochondrial proteins retrieved in our study might have associated with viral RNA prior to mitochondrial import. In this regard, *Hyalomma* orthologues of the thirteen proteins encoded by mitochondrial DNA were not identified among the proteins associated with viral RNA. Alternatively, the mitochondrial proteins associated with the genomic S fragment could have been liberated from damaged mitochondria, which are vulnerable to different types of cellular stress. In a recent study, Jakob *et al.* (32) found that mitochondrial stress induced by infection with the Yellow Fever virus, but not chemically-triggered mitochondrial stress, induced the formation of stress granules that were highly enriched in mitochondrial proteins.

It is tempting to speculate that mitochondrial proteins might be purposefully sequestered by viral RNP. In mammalian cells, such sequestration might antagonize the antiviral function provided by mitochondria. It is also possible that certain mitochondrial proteins are hijacked to the benefit of the virus. The number and diversity of associated proteins would seem to preclude all of the proteins playing a specific role in viral replication, but presumably only a small number of the mitochondrial proteins would be bound directly to the viral RNA or viral proteins N and L: the others having been retrieved within protein complexes. Alternatively, association with mitochondrial proteins might represent a dead-end for the virus. As a result of the host cell immune response, non-replicative viral RNA might be sequestered in cytoplasmic RNA particles, such as stress granules, in association with mitochondrial proteins released from mitochondria as a result of infection-related mitochondrial stress.

In conclusion, we have resolved the first RNA interactome between orthonairovirus RNA and a tick cell proteome. Viral RNA-associated proteins belonged to diverse functional classes and, unexpectedly, a majority were of mitochondrial origin. Whether these proteins represent viral restriction or dependency factors remains to be determined in future functional studies, and critically, whether these results can be transposed to CCHFV, a virus of great societal concern, must await experiments performed in NSB-4 facilities.

## Acknowledgements

This study was supported by Animal Health Department of INRAE. The PhD of ST was financed by a grant from DIM1Health2.0 and the Agence de l’Innovation de Défense. AWC received postdoctoral funding from the LabEx Integrative Biology of Emerging Infectious Diseases.

