## Supplementary File 1 for "Molecular dialogue between Orthonairovirus and tick: RNA-protein interactome of Hazara virus, a BSL2 model of Crimean-Congo Hemorrhagic Fever virus, in *Hyalomma* cells"

|  | **Séquence 5’ 3’** |
| --- | --- |
| HAZV 1 | TGCGGCAACGATATCTTTGA[BIOTEG] |
| HAZV 2 | TAAAGGCAATGCCACCAACA[BIOTEG] |
| HAZV 3 | GGCAGTCCTCAACTATAAGA[BIOTEG] |
| HAZV 4 | GCTCATTGAACTGTTTGCTG[BIOTEG] |
| HAZV 5 | CTACTACTGGCTTTGGAAGG[BIOTEG] |
| HAZV 6 | TAGCAAAGCTGGTTGAGCTA[BIOTEG] |
| HAZV 7 | ATTCGACGCAGGAACAGGAT[BIOTEG] |
| HAZV 8 | GGATGCCAACTACCAAAAGC[BIOTEG] |
| HAZV 9 | ATCCTTTTGTGCCGAAATTC[BIOTEG] |
| HAZV 10 | CCTTTTGAGAGCAAAACCGG[BIOTEG] |
