## Supplementary File 2 for "Molecular dialogue between Orthonairovirus and tick: RNA-protein interactome of Hazara virus, a BSL2 model of Crimean-Congo Hemorrhagic Fever virus, in *Hyalomma* cells"

```
library(data.table)
library(dplyr)
library(curl)


#PARAMETERS####
#USED WHEN EXECUTING BLASTP

threads <- 10 #number of processing units allowed for the execution
evalue <- 1e-3 #maximal evalue filter
max_hits <- 15 #maximum blast hits
min_pident <- 30 #minimal query identity threshold
min_coverage <- 60 #minimal query coverage threshold

#BLAST PATHS####

blastp_path    <- '"C:/Program Files/NCBI/blast-2.17.0+/bin/blastp.exe"'
blastdbcmd_path<- '"C:/Program Files/NCBI/blast-2.17.0+/bin/blastdbcmd.exe"'

#LOCAL DATABASES CREATION####
#(DB MUST HAVE BEEN CREATED LOCALLY FROM PROTEOME FASTA FILES USING -
makeblastdb IN TERMINAL PRIOR TO SCRIPT EXECUTION)

blast_db <- list(
  human      = "C:/blast_db/human/human",
  ixodes     = "C:/blast_db/ixodes/ixodes",
  drosophila  = "C:/blast_db/drosophila/drosophila",
  aedes      = "C:/blast_db/aedes/aedes",
  hyalomma    = "C:/blast_db/hyalomma/hyalomma"
)

uniprot_annotation <- fread("all_species_uniprot_annotation.tsv", header = TRUE, sep =
"\t", stringsAsFactors = FALSE)


#INTEGRITY VERIFICATION OF THE LOCAL DATABASES####

for (db in blast_db) {
  files <- list.files(dirname(db), pattern = paste0(basename(db), "\\.(psq|pin|phr)$"))
  if (length(files) == 0) stop("BLAST db not found : ", db)
}


#CREATING THE FOLDER CONTANING THE BLAST RESULTS####

if (dir.exists("blast_results_name")) unlink("blast_results_name", recursive = TRUE)
dir.create("blast_results_name", showWarnings = FALSE)
```

```
#CREATING THE QUERY FROM AN EXISTING FASTA FILE####

fasta_query <- "ChIRP.fasta"
if (!file.exists(fasta_query) || file.info(fasta_query)$size == 0)
  stop("FASTA file not found or empty : ", fasta_query)

#ID EXTRACTION FROM THE QUERY FASTA FILE####

get_ids_from_fasta <- function(fasta) {
  headers <- readLines(fasta)
  headers <- headers[grepl("^>", headers)]
  ids <- sub("^>(?:sp\\||tr\\|)?([^| ]+).*", "\\1", headers)
  unique(ids)
}

#"unspecified_product" RETRIEVAL####

get_query_annotation_from_fasta <- function(fasta) {
  lines <- readLines(fasta)
  headers <- lines[grepl("^>", lines)]

  data.frame(
    vectorbase_id_source = sub("^>(?:sp\\||tr\\|)?([^| ]+).*", "\\1", headers),
    is_unspecified_product = grepl("gene_product=unspecified product", headers),
    name_prot_hyalomma = sub(".*transcript_product=([^|]+).*", "\\1", headers),
    stringsAsFactors = FALSE
  )
}
query_annotation <- get_query_annotation_from_fasta(fasta_query)


#PRESENCE VERIFICATION OF ALL THE QUERY SEQUENCES IN THE LOCAL DATABASE####

check_sequences_in_db <- function(fasta_query, db_path, blastdbcmd_path){
  message("Presence verification of ", fasta_query, " in ", db_path , " db")
  query_ids <- get_ids_from_fasta(fasta_query)
  chunk_size <- 200
  ids_clean <- unique(query_ids)
  missing_ids <- c()
  chunks <- split(ids_clean, ceiling(seq_along(ids_clean)/chunk_size))
  for(i in seq_along(chunks)){
    id_list <- paste(chunks[[i]], collapse = ",")
    tmp_out <- tempfile(fileext = ".fasta")
    cmd <- sprintf('%s -db "%s" -entry "%s" -out "%s"', blastdbcmd_path, db_path, id_list,
tmp_out)
    system(cmd)
```

```
    if(!file.exists(tmp_out) || file.info(tmp_out)$size == 0){
      missing_ids <- c(missing_ids, chunks[[i]])
    } else {
      extracted_ids <- get_ids_from_fasta(tmp_out)
      missing_in_chunk <- setdiff(chunks[[i]], extracted_ids)
      if(length(missing_in_chunk) > 0) missing_ids <- c(missing_ids, missing_in_chunk)
      file.remove(tmp_out)
    }
  }
  if(length(missing_ids) > 0){
    warning("Missing sequences in the database : ", paste(missing_ids, collapse = ", "))
    return(FALSE)
  }
  message("All query sequences are present in the database\n")
  return(TRUE)
}

if(!check_sequences_in_db(fasta_query, blast_db$hyalomma, blastdbcmd_path)) {
  stop("Error : all query sequences are not found in the database ; please correct before
continuing\n")
}


#BLASTP FUNCTION####

run_blast <- function(query, db, out_file){
  cmd <- sprintf(
    '%s -query "%s" -db "%s" -out "%s" -outfmt "6 qseqid sseqid pident length qlen slen qstart
qend sstart send nident evalue bitscore" -evalue %g -max_target_seqs %d -num_threads
%d',
    blastp_path, query, db, out_file, evalue, max_hits, threads
  )
  message("BLAST execution : ", out_file)
  ret <- system(cmd, intern = FALSE, ignore.stderr = FALSE)
  if(!file.exists(out_file) || file.info(out_file)$size == 0)
    stop("BLAST did not produce valid results ", out_file)
}


#ID EXTRACTION FROM THE DATABASES

get_fasta_from_db <- function(ids, db, out_fasta, chunk_size = 200){
  ids_clean <- unique(sub("^(?:sp\\||tr\\|)?([^|-]+).*", "\\1", ids))
  if(length(ids_clean) == 0) stop("No ID for fasta extraction")
  if(file.exists(out_fasta)) file.remove(out_fasta)
  chunks <- split(ids_clean, ceiling(seq_along(ids_clean)/chunk_size))
  for(i in seq_along(chunks)){
```

```
    id_list <- paste(chunks[[i]], collapse = ",")
    tmp_out <- tempfile(fileext = ".fasta")
    cmd <- sprintf('%s -db "%s" -entry "%s" -out "%s"', blastdbcmd_path, db, id_list, tmp_out)
    system(cmd)
    if(file.exists(tmp_out) && file.info(tmp_out)$size > 0){
      # append to main out_fasta
      cat(readLines(tmp_out), file = out_fasta, sep = "\n", append = TRUE)
      file.remove(tmp_out)
    } else {
      warning("blastdbcmd did not return anything for (IDs: ", substr(id_list,1,200), "...)")
    }
  }
  if(!file.exists(out_fasta) || file.info(out_fasta)$size == 0)
    stop("Empty reverse FASTA from ", db)
  return(out_fasta)
}


#ADJUSTED P% IDENTITY CALCULATION####

compute_adjusted_identity_for_pair <- function(hsps_df, qlen) {
  pos_best_pident <- rep(NA_real_, qlen)
  for(i in seq_len(nrow(hsps_df))){
    qs <- as.integer(hsps_df$qstart[i]); qe <- as.integer(hsps_df$qend[i])
    if(is.na(qs) || is.na(qe)) next
    if(qs < 1) qs <- 1
    if(qe > qlen) qe <- qlen
    if(qs > qe) next
    p <- as.numeric(hsps_df$pident[i])
    idx <- qs:qe
    current <- pos_best_pident[idx]
    to_replace <- is.na(current) | (p > current)
    pos_best_pident[idx[to_replace]] <- p
  }
  covered <- which(!is.na(pos_best_pident))
  union_length <- length(covered)
  if(union_length == 0) return(list(adjusted_pident = NA_real_, adjusted_coverage = 0))
  adjusted_nident_est <- sum(pos_best_pident[covered] / 100)
  adjusted_pident <- (adjusted_nident_est / union_length) * 100
  adjusted_coverage <- (union_length / qlen) * 100
  list(adjusted_pident = adjusted_pident, adjusted_coverage = adjusted_coverage,
union_length = union_length)
}

read_blast_adjusted <- function(path){
  dt <- fread(path, header = FALSE)
  if(ncol(dt) == 0) return(NULL)
```

```
  colnames(dt) <- c("qseqid","sseqid","pident","length","qlen","slen",
                  "qstart","qend","sstart","send","nident","evalue","bitscore")
  dt[,
c("pident","length","qlen","slen","qstart","qend","sstart","send","nident","evalue","bitscore
") := lapply(.SD, as.numeric),
     .SDcols =
c("pident","length","qlen","slen","qstart","qend","sstart","send","nident","evalue","bitscore
")]
  pairs <- unique(dt[, .(qseqid, sseqid)])
  out_list <- vector("list", nrow(pairs))
  for(i in seq_len(nrow(pairs))){
    qid <- pairs$qseqid[i]; sid <- pairs$sseqid[i]
    hsps <- dt[qseqid == qid & sseqid == sid]
    qlen <- unique(hsps$qlen)
    if(length(qlen) != 1 || is.na(qlen)) qlen <- max(hs$qend, na.rm = TRUE)
    adj <- compute_adjusted_identity_for_pair(hsps, qlen)
    best_evalue <- min(hsps$evalue, na.rm = TRUE)
    out_list[[i]] <- data.frame(
      qseqid = qid,
      sseqid = sid,
      adjusted_pident = adj$adjusted_pident,
      adjusted_coverage = adj$adjusted_coverage,
      bitscore = max(hsps$bitscore, na.rm = TRUE),
      evalue = best_evalue,
      sseqid_canonical = sub("^(?:sp\\||tr\\|)?([^|]+).*$", "\\1", sid),
      stringsAsFactors = FALSE
    )
  }
  bind_rows(out_list)
}

#ACCOUNTING FOR INPARALOGS CLUSTERING####
#(INPARALOGS MUST HAVE BEEN SEARCHED FOR USING THE INPARALOGS IDENTIFICATION
SCRIPT)

inparalogues <- fread("inparalogues_hyalomma.csv") #INPARALOGS SEARCH RESULTS
inparalogues[, seqid_canonical := sub("^(?:sp\\||tr\\|)([^|]+).*", "\\1", seqid)]

add_cluster_info <- function(df, inparalogues){
  df <- df %>%
    mutate(sseqid_canonical = sub("^(?:sp\\||tr\\|)([^|]+).*", "\\1", sseqid)) %>%
    left_join(inparalogues[, .(seqid_canonical, cluster_id)],
           by = c("sseqid_canonical" = "seqid_canonical"))
  return(df)
}

#RBH ADJUSTING TO ACCOUNT FOR INPARALOGS
```

```
compute_rbh_with_clusters <- function(forward_df, reverse_df){

  if(is.null(forward_df) || is.null(reverse_df)) return(NULL)

  forward_best <- forward_df %>%
    group_by(qseqid) %>%
    slice_max(bitscore, n = 1) %>%
    ungroup()

  reverse_best <- reverse_df %>%
    group_by(qseqid) %>%
    slice_max(bitscore, n = 1) %>%
    ungroup()

  colnames(forward_best) <- paste0(colnames(forward_best), "_fw")
  colnames(reverse_best) <- paste0(colnames(reverse_best), "_rv")

  join_df <- left_join(
    forward_best,
    reverse_best,
    by = c("sseqid_canonical_fw" = "qseqid_rv")
  )

  join_df <- join_df %>%
    mutate(
      same_cluster = (!is.na(cluster_id_fw) & !is.na(cluster_id_rv) & cluster_id_fw ==
cluster_id_rv),
      is_RBH = (qseqid_fw == sseqid_rv) | same_cluster
    ) %>%
    rename(
      qseqid = qseqid_fw,
      sseqid = sseqid_fw,
      sseqid_canonical = sseqid_canonical_fw,
      adjusted_pident = adjusted_pident_fw,
      adjusted_coverage = adjusted_coverage_fw,
      bitscore = bitscore_fw,
      evalue = evalue_fw,
      cluster_id = cluster_id_fw
    ) %>%
    select(qseqid, sseqid, sseqid_canonical, cluster_id,
         adjusted_pident, adjusted_coverage, bitscore,
         evalue, is_RBH, same_cluster)

  return(join_df)
}
```

```
#BLAST LOOP FOR EACH DATABASE IN blast_db####

results <- list()
species_list <- names(blast_db)

for(sp in species_list){
  message("\n=== Processing species: ", sp, " ===")

  #FORWARD BLAST: QUERY FASTA -> TARGET DB
  fwd_out <- file.path("blast_results_name", paste0("ChIRP_vs_", sp, ".tsv"))
  run_blast(fasta_query, blast_db[[sp]], fwd_out)
  fwd_hits <- read_blast_adjusted(fwd_out)
  if(is.null(fwd_hits) || nrow(fwd_hits) == 0){
    message("Aucun hit forward pour ", sp)
    next
  }
  #ADDING INPARALOGS CLUSTER INFO TO FORWARD HITS
  fwd_hits <- add_cluster_info(fwd_hits, inparalogues)

  #ID EXTRACTION FOR REVERSE BLAST
  target_ids <- unique(fwd_hits$sseqid_canonical)
  rev_query_fasta <- file.path("blast_results_name", paste0("reverse_query_", sp, ".fasta"))
  get_fasta_from_db(target_ids, blast_db[[sp]], rev_query_fasta)

  #REVERSE BLAST: HIT SEQUENCES -> HYALOMMA DB TO CALCULATE RBH
  rev_out <- file.path("blast_results_name", paste0("reverse_", sp, "_vs_hyalomma.tsv"))
  run_blast(rev_query_fasta, blast_db$hyalomma, rev_out)
  rev_hits <- read_blast_adjusted(rev_out)
  if(is.null(rev_hits) || nrow(rev_hits) == 0){
    message("No reverse hit for ", sp)
    next
  }
  #ADDING INPARALOGS CLUSTER INFO TO FORWARD HITS TO ADJUST RBH IF NEEDED
  rev_hits <- add_cluster_info(rev_hits, inparalogues)

  #ADJUSTED RBH CALCULATION
  df_rbh <- compute_rbh_with_clusters(fwd_hits, rev_hits)
  if(is.null(df_rbh) || nrow(df_rbh) == 0){
    message("RBH none for ", sp)
    next
  }
  df_rbh$species <- sp
  results[[sp]] <- df_rbh
}
```

```
#RESULTS SAVING####

summary_all <- bind_rows(results)
if(nrow(summary_all) == 0) stop("No result for all species")
write.table(summary_all, file =
"blast_results_name/ChIRP_vs_all_species_adjusted_RBH_inparalogue_name.tsv",
            sep = "\t", quote = FALSE, row.names = FALSE)

#ORTHOLOGS FILTERING
orthologues <- summary_all %>%
  filter(adjusted_pident >= min_pident,
         adjusted_coverage >= min_coverage,
         is_RBH == TRUE) %>%
  mutate(
    uniprot_id = sub("^(?:sp\\||tr\\|)?([^|]+).*", "\\1", sseqid_canonical),
    vectorbase_id_source = sub("^(?:sp\\||tr\\|)?([^|]+).*", "\\1", qseqid)
    ) %>%
  left_join(uniprot_annotation, by = c("uniprot_id" = "accession")) %>%
  rename(name_prot_hit = protein_name) %>%
  left_join(query_annotation, by = "vectorbase_id_source") %>%
  select(
    qseqid,
    is_unspecified_product,
    sseqid,
    sseqid_canonical,
    cluster_id,
    same_cluster,
    bitscore,
    adjusted_pident,
    adjusted_coverage,
    evalue,
    is_RBH,
    species,
    uniprot_id,
    vectorbase_id_source,
    name_prot_hyalomma,
    name_prot_hit,
    everything()
  )%>%
  select(-espece
  )

write.table(orthologues, file =
"blast_results_name/ChIRP_orthologues_hyalomma_filtered_adjusted_RBH_inparalogues_n
ame.tsv",
            sep = "\t", quote = FALSE, row.names = FALSE)
```

```
cat("\nAnalysis over\n")
cat("Final results :
blast_results_name/ChIRP_orthologues_hyalomma_filtered_adjusted_RBH_inparalogues_na
me.tsv\n")
```
