## Supplementary File 3 for "Molecular dialogue between Orthonairovirus and tick: RNA-protein interactome of Hazara virus, a BSL2 model of Crimean-Congo Hemorrhagic Fever virus, in *Hyalomma* cells"

```
library(data.table)
library(dplyr)
library(igraph)

#PARAMETERS####
#USED WHEN EXECUTING BLASTP

threads <- 10 #number of processing units allowed for the execution
evalue <- 1e-3 #maximal evalue filter
max_hits <- 15 #maximum blast hits
min_pident <- 30 #minimal query identity threshold
min_coverage <- 60 #minimal query coverage threshold

#BLAST PATHS####
blastp_path    <- '"C:/Program Files/NCBI/blast-2.17.0+/bin/blastp.exe"'
blastdbcmd_path<- '"C:/Program Files/NCBI/blast-2.17.0+/bin/blastdbcmd.exe"'

#LOCAL DATABASE CREATION####
#(DB MUST HAVE BEEN CREATED LOCALLY FROM PROTEOME FASTA FILE USING -makeblastdb
IN TERMINAL PRIOR TO SCRIPT EXECUTION)

blast_db_hyalomma <- "C:/blast_db/hyalomma/hyalomma"

#CREATING THE FOLDER CONTANING THE BLAST RESULTS####

output_dir <- "inparalogues_results"
if (dir.exists(output_dir)) unlink(output_dir, recursive = TRUE)
dir.create(output_dir)

#AUTO-BLASTP HYALOMMA VS HYALOMMA####

auto_blast_out <- file.path(output_dir, "hyalomma_vs_hyalomma.tsv")

cmd_blastp <- sprintf('%s -query "%s" -db "%s" -out "%s" -outfmt "6 qseqid sseqid pident
length qlen slen qstart qend sstart send evalue bitscore" -evalue %g -num_threads %d',
                      blastp_path, paste0(blast_db_hyalomma, ".fasta"), blast_db_hyalomma,
auto_blast_out, evalue, threads)

message("AUTO-BLASTP hyalomma vs hyalomma in progress\n")
system(cmd_blastp)

if(!file.exists(auto_blast_out) || file.info(auto_blast_out)$size == 0) {
  stop("AUTO-BLASTP did not return valid results\n")
}

#POTENTIAL INPARALOGS FILTERING####
blast_dt <- fread(auto_blast_out, header=FALSE)
```

```
colnames(blast_dt) <-
c("qseqid","sseqid","pident","length","qlen","slen","qstart","qend","sstart","send","evalue",
"bitscore")

#AUTO-HIT EXCLUSION (100% identity, same start/end)
blast_dt <- blast_dt %>%
  filter(!(qseqid == sseqid & pident == 100 & qstart == sstart & qend == send))

#QUERY COVERAGE CALCULATION
blast_dt <- blast_dt %>%
  mutate(qcov = 100 * length / qlen)

#FILTERING INPARALOGS CANDIDATES
inparalogues <- blast_dt %>%
  filter(pident >= min_pident, qcov >= min_coverage) %>%
  arrange(desc(pident), desc(qcov), desc(bitscore))

if(nrow(inparalogues) == 0){
  message("No inparalogs detected with used parameters\n")
} else {
  message("nparalogs detected : ", nrow(inparalogues))
}

#INPARALOGS CLUSTERING####

#GRAPH CONSTRUCTION
edges <- inparalogues %>% select(qseqid, sseqid)
g <- graph_from_data_frame(edges, directed = FALSE)

#CLUSTER DETECTION
clusters <- components(g)

cluster_df <- data.frame(
  seqid = names(clusters$membership),
  cluster_id = clusters$membership,
  stringsAsFactors = FALSE
)

#INPARALOGS METRICS DISPLAY
cluster_sizes <- table(cluster_df$cluster_id)
message("Number of inparalog clusters detected : ", length(cluster_sizes))
message("Cluster sizes distribution :")
print(sort(cluster_sizes, decreasing=TRUE))

#RESULTS SAVING####

write.table(inparalogues, file = file.path(output_dir, "inparalogues_pairs.tsv"),
```

```
    sep = "\t", quote = FALSE, row.names = FALSE)

write.table(cluster_df, file = file.path(output_dir, "inparalogues_hyalomma.tsv"),
    sep = "\t", quote = FALSE, row.names = FALSE)

cat("\nAnalysis over\n")
cat("Final results : inparalogues_hyalomma.tsv\n")
```
